# Rearing *Istocheta aldrichi* (Diptera: Tachinidae) from field-collected Japanese beetle (*Popillia japonica*): 2. Methods to improve overwintering, adult emergence and longevity

**DOI:** 10.64898/2026.02.10.705140

**Authors:** Paul K. Abram, Simon Legault, Josée Doyon, Victoria Makovetski, Jacob H. Miall, Jean-Philippe Parent, Jason Thiessen, Jacques Brodeur

**Author notes:** Corresponding author: Paul K. Abram.

## Abstract

*Istocheta aldrichi* (Diptera: Tachinidae) is a specialist parasitoid of the invasive Japanese beetle, *Popillia japonica* (Coleoptera: Scarabaeidae). Research and releases for biological control depend on field collecting parasitized hosts and rearing the parasitoid through diapause to obtain *I. aldrichi* adults. This study investigated how rearing practices before, during and after the seasonal overwintering period affected the proportion of *I. aldrichi* pupae that emerged as adults, the timing of parasitoid emergence, and their longevity. Increasing cold exposure duration during overwintering increased adult *I. aldrichi* emergence from puparia and reduced development time after transfer to warm conditions. Adult *I. aldrichi* emergence from overwintered puparia depended on interactions between overwintering environment (indoors *vs*. outdoors), spring thermal regime, and the timing of host collection in the previous season. Burying puparia in the soil in late summer/early fall resulted in higher subsequent adult *I. aldrichi* emergence. Manipulating spring temperatures in controlled environments allowed parasitoid emergence to be staggered over several weeks without reducing emergence success. Emergence under outdoor spring conditions was unreliable. Adult longevity was affected by temperature and diet: cooler conditions extended lifespan, honey-water increased longevity relative to pollen alone or honey-water and pollen together. These results provide a foundation to further improve *I. aldrichi* rearing techniques for use in experimental research and applied biological control of *P. japonica*.

## Introduction

Parasitoid flies in the family Tachinidae are important natural enemies of many pest insects worldwide (Stireman et al., 2006; Cingolani et al., 2025). There are important challenges with rearing tachinids for use in biological control; for example, aspects of their biology or that of their hosts can make them difficult or impossible to rear in year-round continuous colonies (Dindo & Grenier, 2014). Species that cannot be readily reared year-round may still be used in biological control programmes; parasitized hosts can be collected directly from wild populations, reared to the adult stage, and released. This strategy is best applied to classical (importation) biological control, which typically consists of releasing a relatively small number of parasitoids over a short period of time (e.g., a few years) to establish self-sustaining populations. To produce tachinids for this purpose, it is important to have basic knowledge of how to best collect them, rear them through important events in their seasonal life cycle (e.g., diapause), schedule their emergence for biological control releases, and maximize adult survival and early egg production after emergence and before release.

*Istocheta aldrichi* (Mesnil) (Diptera: Tachinidae) is a specialist tachinid parasitoid of the Japanese beetle, *Popillia japonica* Newman (Coleoptera: Scarabaeidae). Both species are native to Japan. Beginning in the early 1900s, *P. japonica* became a major invasive pest of a wide variety of plants cultivated for agriculture and ornamental purposes in North America (Clausen et al., 1927; Potter & Held, 2002). Since then, it has continued to expand its range in North America and has recently become established in Europe (Shanovich et al., 2019; Strangi et al., 2024). Following exploration for natural enemies of *P. japonica* in its native range in the early 1900s, *I. aldrichi* was one among several parasitoid species released for classical biological control of *P. japonica* in the northeastern USA starting in the 1920s (Clausen et al., 1927; Fleming, 1968). The parasitoid established and spread northwards (O’Hara, 2014; Gagnon et al., 2023; Brodeur et al., 2024). Starting in 1990, *I. aldrichi* was successfully redistributed to other areas of the USA (e.g., Minnesota, Colorado, North Carolina) (McDonald & Klein, 2023). There was also a failed attempt to establish the parasitoid in the Azores archipelago (Simões & Grenier, 1999). In areas where *I. aldrichi* did establish, its biological control impact on *P. japonica* was considered to be low or uncertain for about 100 years after its release (Fleming, 1968; Potter & Held, 2002). However, recent studies in Québec, Canada (Gagnon et al., 2023; Legault et al., 2024; Lasnier et al., 2025) and Minnesota, USA (Hutchinson, 2023; Hutchinson et al., 2024) documented relatively high parasitism levels of *P. japonica* by *I. aldrichi* (i.e., observed seasonal average percent parasitism sometimes exceeding 20%), resulting in a renewed interest in this parasitoid from both basic and applied perspectives (Pelletier et al., 2023; Stilwell et al., 2025; Brodeur et al., 2024; Makovetski & Abram, 2024; Makovetski et al., 2025). From 2023–2025 *I. aldrichi* was redistributed from Québec and Ontario to British Columbia (Canada), where *P. japonica* populations were first detected in 2017 (Makovetski & Abram, 2024; CFIA, 2025). The parasitoid is also being considered for introduction to Europe (CABI, 2021), where more recently detected *P. japonica* populations (currently present in Italy and Switzerland) are spreading (Strangi et al., 2024).

Despite the long history of biological control programmes using *I. aldrichi*, relatively little information is available on the best methods to rear these flies in support of experimental biology studies and biological control releases. Maintaining the parasitoid in continuous laboratory colonies has not been attempted, to our knowledge, probably because its host, *P. japonica*, is itself labor-intensive to rear under laboratory conditions (Ludwig & Fox, 1938). Rather, previous *I. aldrichi* introduction programs have relied on collecting parasitized *P. japonica* from natural populations, using hand collections or traps with olfactory attractants (Clausen et al., 1927; McDonald & Klein 2023). Parasitized beetles are easy to identify because the egg of *I. aldrichi*, typically laid on the host pronotum, is visually obvious. The parasitized beetles are then reared on foliage on top of a substrate (e.g., soil, sphagnum moss) in which they bury themselves before they die and the parasitoid larva forms a puparium (e.g., Clausen et al., 1927; Fleming, 1968). Initial releases in the USA sourced *I. aldrichi* from parasitized *P. japonica* cadavers shipped from Japan. These cadavers (and associated *I. aldrichi* puparia) were held outdoors under the soil or in indoor growth chambers adjusted to match outdoor soil temperatures, with the highest emergence of *I. aldrichi* adults observed when puparia were overwintered outdoors (King & Hallock, 1925; Fleming, 1968). Later experiments done in the Azores archipelago (Simões & Grenier, 1999) suggested that *I. aldrichi* required a 1–4 month period of chilling (at 4°C in a refrigerator) to complete their diapause and emerge as adults, with emergence being the highest for the longest chilling periods. Other practitioners rearing *I. aldrichi* for redistribution reportedly overwintered puparia in containers buried outdoors in soil until the spring, whereupon puparia were unearthed and left at release sites in protected cages until adult emergence (McDonald & Klein, 2023). Adult *I. aldrichi* are reported to feed on plant nectar (McDonald & Klein, 2023) but the effect on adult longevity and fecundity, and whether other potential sources of nutrition (e.g., pollen) might be important, is not known. While these past, mostly descriptive reports of *I. aldrichi* rearing provided a good foundation for contemporary biological control efforts, there is still a need to quantify what factors may be important to optimize emergence of *I. aldrichi* from field-collected parasitized *P. japonica*. Varying the parasitoid’s spring rearing conditions to schedule their emergence so that biological control releases occur during periods of high *P. japonica* abundance is one particular aspect that has not been adequately explored.

In this study, the importance of the following factors affecting *I. aldrichi* adult emergence success were tested experimentally: (i) length of time the puparia extracted from parasitized beetles were held in cool conditions; (ii) time of year when *P. japonica* adults were collected in the field the previous season; (iii) overwintering conditions (i.e., indoors versus outdoors under the soil, burial depth and timing); (iv) post-overwintering temperature in the spring. In addition, to test how the longevity of *I. aldrichi* could be extended between emergence and release of adults in biological control programmes, (v) different combinations of diet and storage temperature on adult survival were tested. These methods were tested and applied from 2022–2023 as part of a redistribution programme for *I. aldrichi* from Ontario and Québec to British Columbia (Canada) (Makovetski & Abram, 2024; Abram, 2026). The effect of various factors on the upstream stages of *I. aldrichi* rearing – that is, how parasitized *P. japonica* are collected and fed before *I. aldrichi* puparia are extracted – were tested in a companion study (Legault et al., 2026).

## Materials and Methods

For clarity, the series of experiments described below is numbered based on the life cycle progression of *I. aldrichi* rather than the true chronological order in which they were conducted.

### Collections of parasitized *P. japonica*

For Experiments 1, 2 and 4, parasitized *P. japonica* were collected in the Ottawa (Ontario, Canada) and Saint-Jean-sur-Richelieu (Québec, Canada) regions between July 4 and August 3, 2022. Adult *P. japonica* were hand-collected or caught in commercially available traps baited with dual-lure food and sex pheromone attractants (Legault et al., 2024). Live parasitized *P. japonica* with at least one macrotype parasitoid egg, characteristic of *I. aldrichi*, were placed in cohorts (n = 77) of 43 to 100 beetles (total n = 6,211 beetles) with each cohort collected on the same day. Each cohort of parasitized beetles were placed in rearing chambers consisting of plant pots (15 × 17.5 cm, ITML, Ontario, Canada) containing standard potting soil or vermiculite with a fine-mesh sleeve supported by two metal hoops (wire coat hangers) or a plastic stake placed over the top. In one laboratory (Saint-Jean-sur-Richelieu), additional cohorts (n = 8) of unknown numbers of *P. japonica* collected on the same day were reared in larger containers to provide additional *I. aldrichi* puparia (total n = 1,009 puparia) for experiments. These large containers (61 L, 60.3 × 40.6 × 34.3 cm) were made of clear plastic, were ventilated with meshed openings on the top and sides, and contained an 8 cm high layer of moist vermiculite. Foliage from known host plants, *Vitis riparia* (Michaux) (Vitales: Vitaceae) and *Parthenocissus quinquefolia* (L.) Planchon (Vitales: Vitaceae), collected from outdoors was provided *ad libitum* every 1–3 days for 7–14 days to feed the adult *P. japonica*; after this period most beetles have died from parasitism (J. Miall, J.-P. Parent, S. Legault, J. Doyon, personal observations; Clausen et al., 1927). Rearing conditions during this period were 21–24 °C, 40–70% RH, and 16:8 light: dark cycles. In the fall (late September and early October, 2022), dry foliage was removed from samples and the substrate containing *P. japonica* cadavers and *I. aldrichi* puparia was transferred to 14.0–15.5°C in the dark. The samples were then shipped to Agassiz, British Columbia, Canada (October 20–November 1, 2022) to manually extract *I. aldrichi* puparia from *P. japonica* cadavers, as well as those in the soil that had fallen out of degraded *P. japonica* specimens; approximately 25–50% of *I. aldrichi* larvae exit the beetle cadaver themselves before pupation (S. Legault and J. Doyon, personal observations). For samples where the initial number of *P. japonica* was known (n = 77), the proportion of parasitized beetles yielding *I. aldrichi* puparia was therefore approximated as the number of puparia extracted from a sample divided by the initial number of beetles placed in the sample. On November 10, 2022, puparia were placed in cohorts of up to 100 in labeled Solo cups (2 oz) containing moistened vermiculite, and stored in a walk-in growth chamber at 5°C and 80% RH in darkness and misted with water as needed until use in experiments. The effect of superparasitism was not accounted for; i.e., the fact that some beetles had more than one *I. aldrichi* egg on them. Superparasitism may have had modest downstream effects on the quality of surviving puparia in superparasitized hosts (J. Doyon, S. Legault and J. Brodeur, unpublished data).

For Experiment 3, parasitized beetles were arbitrarily hand-picked in 2024, from July 12 to 17 at a vineyard in Saint-Paul-Abbotsford, Québec. *Popillia japonica* adults bearing at least one *I. aldrichi* egg on their pronotum were placed in a container with fresh leaves (*Vitis* spp.) and brought back to the laboratory. They were then placed in cohorts of up to 200 individuals in 1L ventilated plastic containers with moistened vermiculite for substrate. Containers were stored in a controlled environment (21 ± 1°C, 45 ± 5 % RH, 14L: 10D), and parasitized beetles were fed *ad libitum* with fresh vine leaves for two weeks. After that period, the substrate of each container was spread on a sorting tray and carefully examined to retrieve beetle cadavers and loose *I. aldrichi* pupae. Beetle cadavers were dissected under a stereomicroscope to extract all remaining *I. aldrichi* pupae. Parasitoid puparia were stored at 21□±□1 °C, 45□±□5 % RH, 14L: 10D in 2 oz plastic cups filled with 0.5 cm of moist vermiculite until their use in the experiment.

Representative specimens of *I. aldrichi* were verified by Dr. James O’Hara (Canadian National Collection of Arthropods and Nematodes [CNC], Agriculture and Agri-Food Canada, Ottawa, Canada) and vouchers are incorporated into the CNC.

### Experiment #1: Effects of overwintering duration on *I. aldrichi* emergence and development time

The goals of the first experiment were to determine: (i) how long *I. aldrichi* puparia must be kept at cool temperatures in order to complete their diapause and subsequently develop and emerge as adults at warmer temperatures (as indicated by the proportion of puparia emerging as adults); and (ii) how long it takes *I. aldrichi* adults to emerge from puparia at warm temperatures after cold exposure of varying durations. Three to four randomly selected samples of between 13 and 93 puparia (subsampled from collections from July 17–19, 2022 from Saint-Jean-sur-Richelieu), were removed at each of six intervals (51, 75, 97, 119, 139, and 163 days) after the puparia (which were of similar initial ages because of similar collection dates) were first placed in a growth chamber set to 5°C (actual average from a temperature logger: 5.51°C). These samples (total n = 19 samples, n = 736 puparia) were then incubated in a growth chamber at 22 ± 1°C, 16:8h L:D, and 40-60% RH. Adult *I. aldrichi* emergence was checked every 2-3 days. Proportion emergence was calculated as the number of emerged adult *I. aldrichi* divided by the number of puparia in each sample. Development time was calculated as the number of days since the sample was placed at 22°C until emergence, which was designated as the midpoint between the last time emergence was checked and the time the individual was observed. To avoid pseudoreplication, the average time until emergence of *I. aldrichi* in each sample was used in the subsequent statistical analysis.

### Experiment #2: Effects of overwintering and spring conditions on survival and development time of *I. aldrichi*

The second experiment tested the effects of winter and spring temperature conditions on the level and timing of *I. aldrichi* emergence in the summer. More specifically, whether: (i) overwintering *I. aldrichi* puparia indoors at a constant temperature produced comparable emergence levels to overwintering them under the soil outdoors; and (ii) keeping puparia at constant low temperatures then immediately bringing them to room temperature produced similar emergence levels to gradually increasing the temperature of puparia in the spring (in the field or in incubators). The treatments were not designed to isolate individual factors affecting fly emergence and development; rather, they were intended to compare different *I. aldrichi* rearing ‘systems’ comprising a suite of possible practices that could plausibly be used in biological control programmes.

From a subset of the initial set of puparia samples (see above), 15 cohorts of puparia (seven from Saint-Jean-sur-Richelieu and eight from Ottawa) were created, with each cohort of 118–985 puparia (total n = 4,329) collected in the same region on the same date. Collection dates of the cohorts of puparia varied from July 11 to August 3, 2022. Each group was then divided into eight subsamples (14–123 puparia each) and assigned to one of eight conditions. These conditions were full factorial combinations of two overwintering treatments (storage conditions) between November 10, 2022 and April 6, 2023) and four spring treatments (storage conditions) between April 6 and adult fly emergence). The spring treatment started when weekly outdoor soil temperatures reached ∼10°C, estimated to be near the lower developmental threshold of *I. aldrichi* puparia (P. K. Abram, unpublished analyses). The subsamples were treated as the unit of replication; thus, there were 15 replicates of each of the eight treatments. The non-independence of the subsamples with respect to origin and group was accounted for in the statistical analysis (see *Statistical analyses*).

The two overwintering treatments were ‘indoors’ and ‘outdoors’. Subsamples of puparia overwintered indoors were placed in 300 mL of moist vermiculite/soil mix (1:1) in fine-mesh bags (9 × 18 cm) in a walk-in growth chamber at 5°C. Puparia overwintered outdoors were placed in the same bags and soil mix but in order to protect them from wildlife they were placed inside metal wire mesh bird feeders (14.5 × 14.5 × 14.5 cm; mesh size: 1 × 0.5 cm, NO/NO feeders, Amazon, Canada) and buried ∼20 cm below the soil surface in a research plot in Agassiz, British Columbia (GPS: 49.240, −121.754). At this soil depth, average temperatures (mean: 3.45°C; minimum: −1.08°C; maximum: 9.10°C) are similar to outdoor air temperatures (mean: 3.68°C; minimum: −11.62°C maximum: 12.72°C) but extremes are moderated. The samples were buried under naturally occurring soil which was lightly compacted. A metal grate was also placed on the surface to prevent vertebrate wildlife from digging. Temperatures during overwintering in the two treatments over time, compared to daily average air temperatures, are shown in Figure S1 (Supplementary Material).

The four spring treatments were:

I. Held outdoors (same location as above) in a plastic tote embedded in the soil and covered with white bags of potting soil, designed to produce internal temperatures similar to a temperature probe buried in *in situ* soil at 20 cm depth. Samples were elevated with cardboard to prevent direct heat transfer from the plastic or drowning in any accumulated water.
II. Placed in an incubator that was programmed once weekly to the previous week’s average soil temperature, as determined by a temperature probe (Onset Inc., Hoskin Scientific, Canada) logging every 10 minutes buried 20 cm below the soil surface at the location described above.
III. Same as (II), but in order to delay fly emergence 1.5°C was subtracted from the temperature setpoint each week.
IV. Kept in a growth chamber set to 5°C (actual average: 5.46°C) until June 19, 2023 and then moved to an incubator set to 23.0°C (actual average: 22.76°C).

Temperature readings from each of the four spring treatments are shown in Figure S2 (Supplementary Material). To facilitate collection of emerging adult *I. aldrichi*, puparia were moved to plastic ventilated emergence containers (9.5 × 6 × 6 cm) containing 200 mL of vermiculite/soil mix (1:1) before being transferred to each of the four spring treatments. To prevent emerging *I. aldrichi* adults from dying before they could be counted, liquid honey was smeared on the underside of the container lid *ad libitum* starting when the first fly emerged in each treatment. Fly emergence in each treatment was checked every 2–5 days, with emergence time designated as the midpoint between the last time emergence was checked and the time the individual was observed. Development time was calculated as the number of days elapsing between the start of the spring treatment (April 6) and emergence.

### Experiment #3: Effects of burying date and spring cool storage duration on *I. aldrichi* emergence

The third experiment tested the effects of the timing of burial of *I. aldrichi* puparia under the soil in the late summer/fall on *I. aldrichi* emergence, when subsequent spring emergence was staggered with temperature as would typically be done for biological control releases (see above). Groups of 25–46 puparia were transferred from standard rearing conditions (see “Collections of parasitized *P. japonica*” above), to small muslin bags (3 × 4 cm), and buried outdoors in the soil of a lawn plot at the Montréal Botanical Garden, Canada (GPS: 45.561, - 73.572), 10–40 cm under the soil on one of three dates (August 30 – ‘Late August’, September 24 – ‘Late September’, or October 28 – ‘Late October’). Puparia were unearthed on May 21, 2025 and puparia from each sample were split randomly in three groups and transferred to ventilated insect rearing boxes (7 × 7 × 10 cm, ProLab Scientific, Laval, Québec, Canada). Lids of rearing boxes were removed, and boxes were transferred to three Bugdorm insect rearing cages (33 × 33 × 77 cm, MegaView Science Co. Ltd., Taichung, Taiwan). Cages were then stored in cool dark conditions (12 ± 1°C, 50 ± 5% RH, 0L: 24D) for one of three durations (37, 47 or 70 days) and then transferred to warmer conditions (22 ± 1°C, 80 ± 5% RH, 16L: 8D). Thus, there were a total of 9 treatments (3 replicates per treatment) representing full factorial combinations of each of the burial date and cool storage duration treatments. After the cool storage periods, cages were monitored daily for adult emergence. After the adult parasitoid emergence period was complete, puparia were examined individually to measure the proportion that yielded adult parasitoids.

### Experiment #4: Effect of storage temperature and nutrition on longevity of *I. aldrichi* adults

The fourth experiment tested which food diets and temperatures could extend the adult life of *I. aldrichi* after emergence. In growth chambers, two temperature treatments at 16:8 h L:D cycles and 40–60% RH were tested: warm (temperature during the 16h light period: 22.0 ± 0.3°C; temperature during the 8h dark period: 20.0 ± 0.3°C) and cool (16.0 ± 0.3°C and 12.0 ± 0.3°C during the light and dark periods, respectively). It was reasoned that the cool temperatures, which are within the range of day and night temperatures that *I. aldrichi* adults could experience in the early summer in Canada, would extend the lives of adult *I. aldrichi*, in line with many past studies of tachinids (e.g., Dindo et al., 2019).

There were three diet treatments: honey (40% honey/water solution w/w), pollen (40% apple pollen/water solution w/w), and honey + pollen (40% honey and 40% apple pollen w/w). These treatments were selected based on a subset of those tested in Dindo et al. (2019). In addition, a preliminary experiment showed that honey and pollen extended the life of *I. aldrichi,* relative to water, at constant 23°C, and that honey-water was preferable to sucrose solution because it produced less unwanted fungal growth. Individual *I. aldrichi* adults, emerging from puparia in Experiment 1 and collected within 48 h of emergence, were assigned sequentially as they emerged to one of the six factorial combinations of the temperature and diet treatments and placed individually in ventilated plastic containers (9.5 × 6 × 6 cm). Male and female *I. aldrichi* adults, whose sex was determined based on antennal and abdominal morphological features upon emergence (J. O’Hara, personal communications), were distributed evenly among treatments as they emerged. After eliminating individuals that died due to experimenter error, the sample sizes of *I. aldrichi* per temperature/diet combination ranged from 7–10 and 9–10 male and female individuals, respectively, for a total of 17–19 individuals per treatment (n = 106 total individuals). At the start of each replicate 300 ml of the assigned diet solution were pipetted onto cellulose vial plugs (diameter: 2.2 cm, height: 2.6 cm height, Diamed, Canada) and these plugs were placed on the bottom of the container and refreshed every 2–3 days. One Kimwipe tissue (Kimtech Inc., Canada) was placed in each container to provide structure and a dry place for the adult *I. aldrichi* to rest. The status of the adults (alive, dead) was checked every 2–3 days, and their time of death was designated as the midpoint between the last two observations. Adult longevity was calculated as the number of days elapsed between the start of the trial and their time of death.

### Statistical analyses

All analyses were conducted in R version 4.4.1 (R Core Team, 2024). Assumptions and quality of fit of all statistical models were verified with residuals and QQ-plots using base R functions as well as the diagnostic functions in the DHARMa package (Hartig, 2024). The significance of pairwise differences between groups was assessed with the R package emmeans (Lenth, 2023) using the Tukey p-value adjustment method.

To test how the duration of cold exposure affected subsequent emergence of *I. aldrichi* in Experiment 1, emergence probability was modeled as a saturating function of chilling duration. A nonlinear asymptotic model was implemented, relating the proportion of puparia that emerged to the number of cold days, with the probability value of the asymptote (p_max_) and half-saturation constant (K) estimated as model parameters. This model was judged to fit the data better than a linear or quadratic model based on model diagnostics. The model was fit by nonlinear least squares, weighted by the total number of puparia per experimental unit. To model how the duration of cold exposure affected the development time of puparia, based on visual inspection of the non-linear relationship and poor fit of other candidate models (i.e., linear, polynomial) emergence timing was modeled as a four-parameter logistic function of cold exposure (Ritz et al., 2015). Here, parameter L represents the expected emergence time at the lowest cold exposure duration, U the emergence time after the longest cold exposure duration, X_50_the cold exposure duration at which emergence timing is halfway between these extremes, and S controls the steepness of the transition. Parameter estimates for the nonlinear model were determined using non-linear least squares. Significance of individual parameters was assessed using t-tests.

To test which factors affected the proportion of *I. aldrichi* puparia that emerged as adults the following season in Experiment 2, a generalized linear mixed model with a binomial error distribution was used. Overwintering treatment, spring treatment, the collection date (numeric calendar date) of the puparia, and their interactions were included as fixed effects. The cohort to which the puparia belonged and their collection origin were included as random effects. To test how the same set of predictors affected the mean development time of puparia, a GLMM with a Gamma error distribution was used. This model did not converge when it contained random effects, whose variance components were estimated at near-zero; therefore, these random effects were removed to improve numerical stability (Barr et al., 2013). Statistical significance of the predictors for both above models was assessed with Type II Wald chi-squared tests.

To test how burial date, the duration of storage under cool conditions, and their interaction affected the proportion of *I. aldrichi* puparia that emerged as adults in Experiment 3, a generalized linear model (GLM) with a binomial error distribution fit with quasi-likelihood was run. Type II F-tests were used to test the statistical significance of each predictor term.

To test what factors affected the longevity of adult *I. aldrichi* in Experiment 4, a Cox proportional hazards model was implemented with temperature treatment, diet treatment, the sex of *I. aldrichi* adults, and their interactions as predictors. Type II Likelihood ratio tests were used to test the statistical significance of each predictor term.

## Results

### Experiment #1: Effects of overwintering duration on *I. aldrichi* emergence and development time

The proportion of *I. aldrichi* puparia emerging as adults depended on how long they were held at 5°C prior to being removed and placed at 22°C to develop. Proportion emergence followed an increasing saturating relationship with cold exposure duration (Table 1, p_max_), with a predicted proportion emergence of 0.54 after 51 days of cold exposure increasing to 0.73 after 163 days, with diminishing returns in proportion emergence with increasing cold exposure duration (Figure 1A).

**Figure 1.**
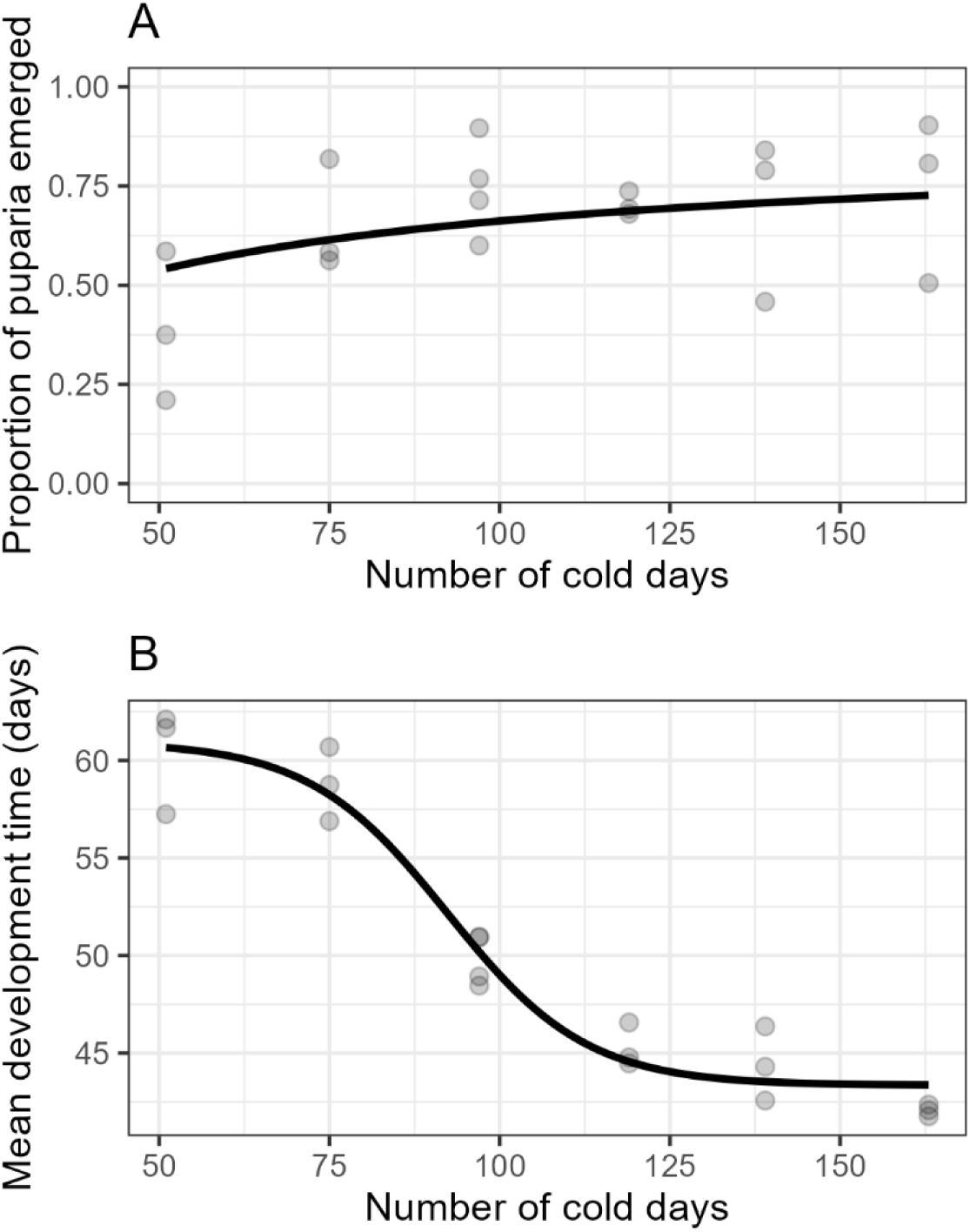
The effect of the duration of puparia cold exposure on: (A) the proportion adult emergence of *Istocheta aldrichi*; and (B) the mean developmental time (i.e., time until adult emergence) of *I. aldrichi* at 22.5°C after being removed from the cold. Black regression lines are from: (A) a non-linear asymptotic model, y = 0.86x/(29.9 + x); (B) a four-parameter log-logistic fit, y = 60.97 - 17.62 / (1 + exp(-(x - 92.35) / 10.23)). Each grey point represents a group of puparia. See Table 2 for statistical information.

**Table 1.** Statistical information for Experiment 1, which tested the effect of the number of days *Istocheta aldrichi* puparia were stored at 5°C on the subsequent proportion adult emergence, and how long emergence took at 22°C. p_max_ – probability value of the asymptote; K – half-saturation constant; L – expected emergence time at the lowest cold exposure duration; U – emergence time after the longest cold exposure duration; x_50_ – cold exposure duration at which emergence timing is halfway between these extremes; S – steepness of the transition. See Figure 1 for model equations and visual representations of model fits to the data.

| Response variable | Model term | Test statistic<sub>df</sub> | p-value |
| --- | --- | --- | --- |
| Proportion adult emergence<sup>a</sup> | $p_{\max}$ | $t_{1,17} = 5.09$ | < 0.0001 |
| | $K$ | $t_{1,17} = 1.17$ | 0.26 |
| Days to adult emergence<sup>b</sup> | $L$ | $t_{1,15} = 47.62$ | < 0.0001 |
| | $U$ | $t_{1,15} = 58.49$ | < 0.0001 |
| | $x_{50}$ | $t_{1,15} = 31.26$ | < 0.0001 |
| | $s$ | $t_{1,15} = 3.53$ | 0.0031 |

<sup>a</sup> Nonlinear asymptotic model.
<sup>b</sup> Four-parameter log-logistic model.

**Table 2.** Statistical information for Experiment 2, which tested the effect of overwintering and spring conditions on the proportion of *Istocheta aldrichi* puparia that emerged as adults and the number of days it took the adults to emerge. See Figures 2 and 3 for visual representations of model fits to the data.

| Response variable | Model term | Test statistic<sub>df</sub> | p-value |
| --- | --- | --- | --- |
| Proportion emergence<sup>a</sup> | Spring treatment (ST) | $\chi^2_{3,86} = 68.10$ | < 0.0001 |
| | Overwintering treatment (OT) | $\chi^2_{1,86} = 19.92$ | < 0.0001 |
| | Collection date (CD) | $\chi^2_{1,86} = 2.26$ | 0.13 |
| | ST × OT | $\chi^2_{3,86} = 50.72$ | < 0.0001 |
| | ST × CD | $\chi^2_{3,86} = 16.39$ | < 0.001 |
| | OT × CD | $\chi^2_{1,86} = 4.23$ | 0.040 |
| | ST × OT × CD | $\chi^2_{3,86} = 10.91$ | 0.012 |
| Days to adult emergence<sup>b</sup> | ST | $\chi^2_{3,72} = 0.38$ | 0.54 |
| | OT | $\chi^2_{1,83} = 587.55$ | < 0.0001 |
| | CD | $\chi^2_{1,83} = 18.16$ | < 0.001 |
| | ST × OT | $\chi^2_{3,72} = 1.70$ | 0.64 |
| | ST × CD | $\chi^2_{3,72} = 0.14$ | 0.71 |
| | OT × CD | $\chi^2_{1,72} = 5.94$ | 0.11 |
| | ST × OT × CD | $\chi^2_{3,72} = 0.06$ | 1.00 |

<sup>a</sup> Generalized linear mixed model with binomial error distribution. Information for random effects terms (collection origin, puparium group; see Methods) are not shown.
<sup>b</sup> Generalized linear model with Gamma error distribution. Test statistics and p-values for significant terms are from a simplified model containing only OT and CD; statistics for the other terms are from the full model containing all terms.

The time required for *I. aldrichi* pupae to emerge as adults after being placed at 22°C (i.e., development time) also depended on how long they were held at 5°C. Time until adult emergence followed a log-logistic relationship with cold exposure duration (Table 1). Predicted development time declined from an upper asymptote of approximately 61 days following shorter cold exposure to a lower asymptote of approximately 43 days after more extended cold exposure (Figure 1B).

### Experiment #2: Effects of overwintering and spring conditions on survival and development time of *I. aldrichi*

Adult emergence of *I. aldrichi* from overwintered puparia depended on interactions between the overwintering environment and spring thermal treatment. These effects also varied with the date on which puparia were collected during the previous season (Table 2; Figure 2). This was reflected by a significant three-way interaction effect among the three variables (Table 2).

**Figure 2.**
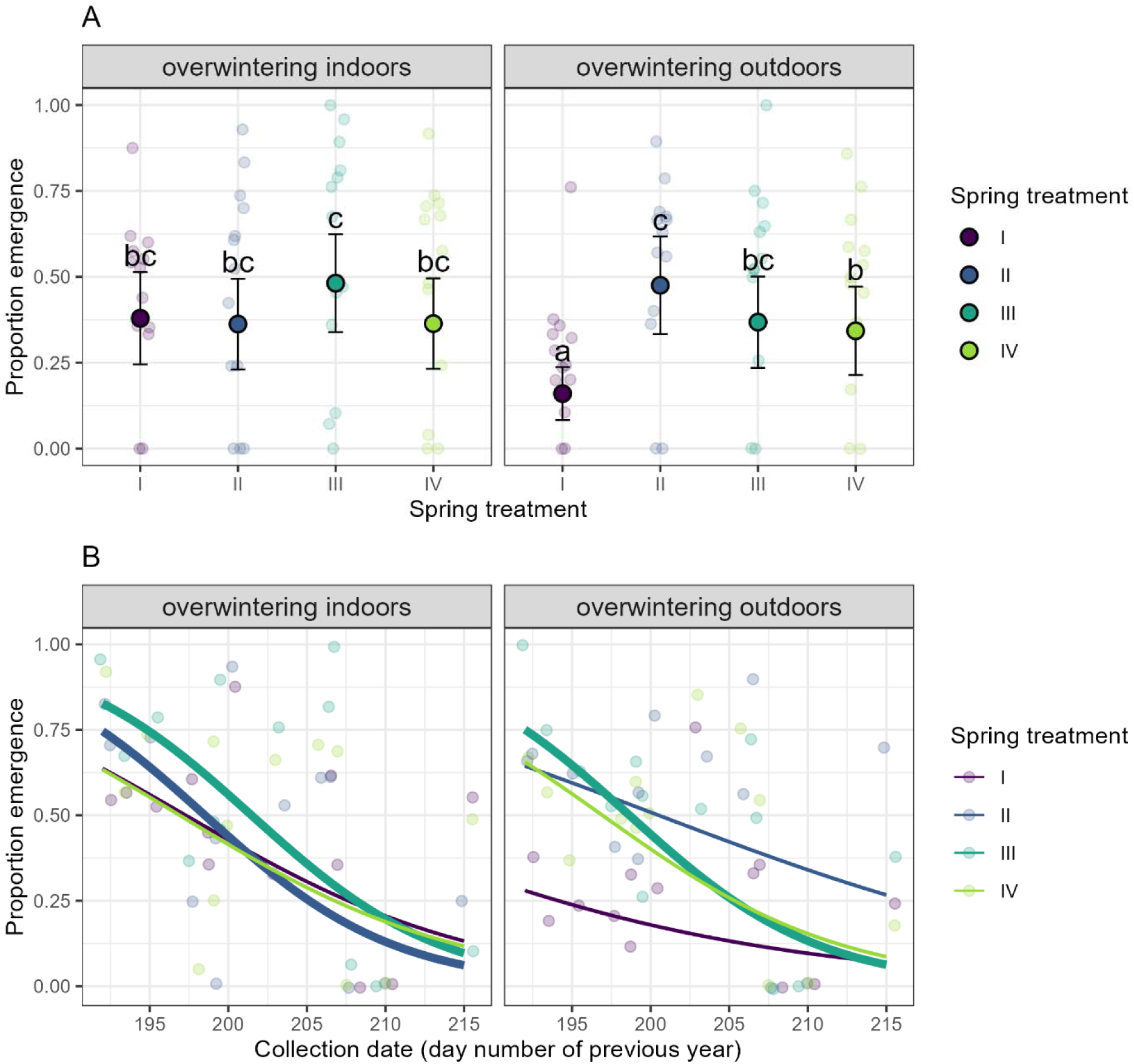
Effect of overwintering (overwintering indoors at 5°C *versus* outdoors at soil temperatures) and spring treatments (I – outdoors under the soil; II – indoors at soil temperatures; III – indoors at lower-than-soil temperatures; IV – indoors at 5°C and then moved to 22°C) on the subsequent proportion emergence of *Istocheta aldrichi* puparia. (A) The estimated marginal mean proportion emergence (black dots, ± 95% CI) in different overwintering and spring treatments, calculated from a GLMM fit; marginal means labeled with different letters are significantly different (Tukey multiple comparisons tests; p < 0.05). (B) The effect of collection date of the parasitized beetles (the prior year) on proportion emergence of puparia in different overwintering and spring treatments. Trend lines are marginal predictions from the fitted GLMM; thicker lines show statistically significant negative trends (Wald *z*-test, p < 0.05). See Table 3 for statistical information.

**Table 3.** Statistical information for Experiment 4, which tested the effect of sex, diet, and temperature on *Istocheta aldrichi* adult longevity. See Figure 5 for model equations and visual representations of model fits to the data.

| Response variable | Model term | Test statistic<sub>df</sub> | p-value |
| --- | --- | --- | --- |
| Longevity<sup>a</sup> | Sex | $\chi^2_1 = 0.75$ | 0.39 |
| | Diet | $\chi^2_2 = 14.61$ | < 0.001 |
| | Temperature | $\chi^2_1 = 9.91$ | < 0.01 |
| | Sex × Diet | $\chi^2_2 = 0.63$ | 0.73 |
| | Sex × Temperature | $\chi^2_1 = 0.0051$ | 0.94 |
| | Diet × Temperature | $\chi^2_2 = 0.59$ | 0.74 |
| | Sex × Diet × Temperature | $\chi^2_2 = 0.27$ | 0.87 |

<sup>a</sup> Cox proportional hazards model. Test statistics and p-values for significant terms are from a simplified model containing only Diet and Temperature. Statistics for the other terms are from the full model containing all terms.

Emergence probabilities differed among spring treatments, but which spring conditions produced the highest and lowest levels of emergence depended on overwintering treatment (Table 2; Figure 2A). For puparia overwintered indoors, predicted proportion emergence was similar (model estimates ranging from 0.36–0.48) for all four treatments. For puparia overwintered outdoors, proportion emergence tended to be highest (0.47) in growth chambers set at outdoor soil temperatures (treatment II) and lowest (0.16) when puparia emerged under outdoor spring conditions (treatment I), with treatments III and IV showing intermediate proportion emergence (0.34–0.37) (Table 2; Figure 2A).

*Istocheta aldrichi* emergence generally declined when samples were collected later in the previous season, but the magnitude of this decline varied depending on both the spring treatment and how insects were overwintered (Table 2; Figure 2B). Declines in proportion emergence with later collection dates tended to be steeper when puparia were overwintered indoors (declining from predicted values of 0.63–0.83 to 0.06–0.13 over the range of collection dates) compared to when they were overwintered outdoors (declining from ranges of 0.27–0.75 to 0.06–0.27) (Figure 2B).

The time until adult *I. aldrichi* emergence was most strongly affected by spring treatment, and increased when samples were collected later in the previous year (Table 2; Figure 3). Development time was not affected by overwintering treatment or interactions among any of the factors (Table 2). The four spring treatments produced *I. aldrichi* emergence that was staggered over a period of 40 days (June 21 to July 31). Predicted average development time per sample was the shortest across the range of collection dates (86.5–90.9 days) when puparia were held in incubators at average spring soil temperatures (spring treatment II) and longest when puparia were held at 5°C until June and then transferred to 23°C (103.7–109.0 days) (spring treatment IV), with intermediate average development times for the other two treatments (Figure 3).

**Figure 3.**
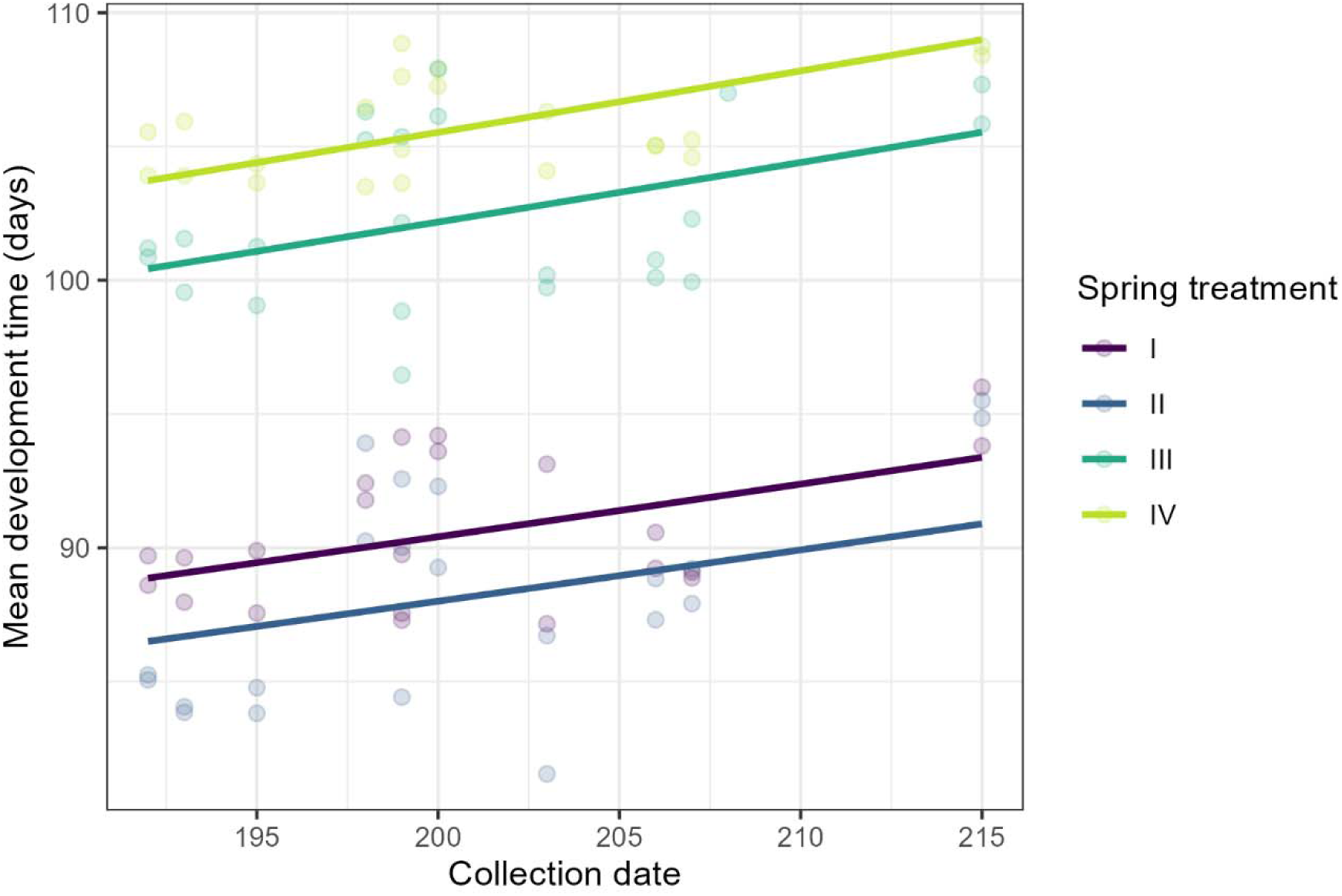
Time until emergence of adult *Istocheta aldrichi* from puparia in different spring treatments (I – outdoors under the soil; II – indoors at soil temperatures; III – indoors at lower-than-soil temperatures; IV – indoors at 5°C and then moved to 22°C) depending on the date (day number of the calendar year) they were collected the previous year. Curves show GLM predictions (see Table 3 for statistical information); light colored points show the mean development time of each independent replicate; overwintering treatments are pooled within spring treatments because overwintering treatment was not a significant predictor of development time.

### Experiment #3: Effects of burying date and spring cool storage duration on *I. aldrichi* emergence

The proportion of *I. aldrichi* emerging was strongly reduced by later burial dates the previous season (F_2,24_ = 151.27, p < 0.0001), with lower emergence at later burial dates (Figure 4). Emergence was approximately 36% when puparia were buried in late August, ∼17% when they were buried in late September, and ∼1% when they were buried in late October. Proportion emergence was similar between spring cool storage durations (F_2,18_ = 3.93, p = 0.14) and there was no effect of the interaction between spring storage duration and burial date (F_4,18_ = 3.93, p = 0.63).

**Figure 4.**
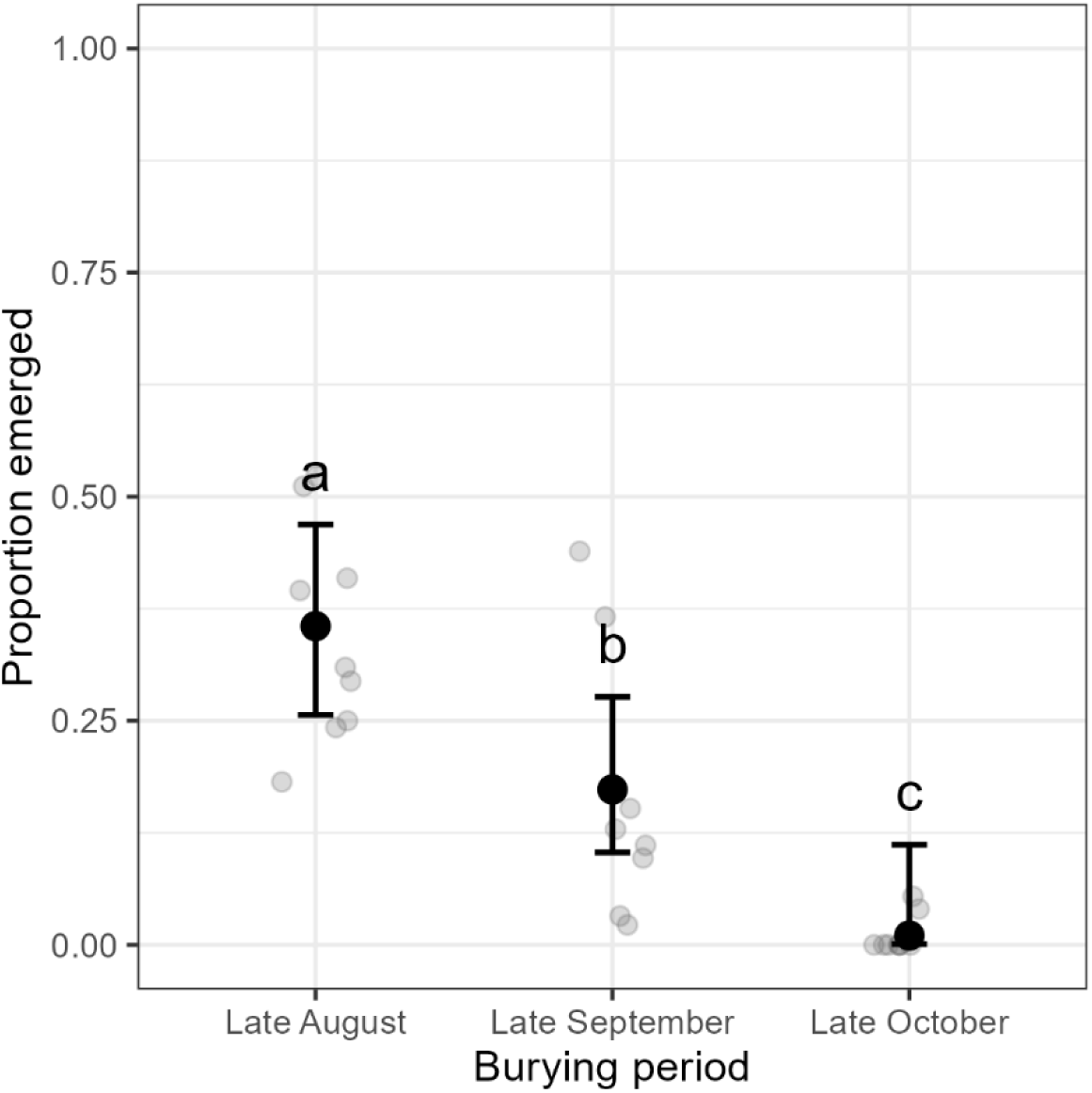
The proportion of *Istocheta aldrichi* adults emerging when puparia were buried at three different periods during the previous season. Each grey symbol shows the proportion yield of a group of 25 to 46 puparia. Black dots show predictions (± 95% CI) from a GLM fitted to the data; marginal means labeled with different letters are significant different (Tukey multiple comparisons tests; p < 0.05).

### Experiment #4: Effect of temperature and nutrition on longevity of *I. aldrichi* adults

Diet and temperature both affected the longevity of adult *I. aldrichi*, and longevity was similar between males and females (Figure 5; Table 3). Adult *I. aldrichi* lived longer under cool temperature conditions than warm temperature conditions (Figure 5). Feeding adult *I. aldrichi* with honey led to the greatest longevity across both temperature treatments, whereas individuals fed honey and pollen or pollen alone had reduced longevity (Figure 5; Table 3).

**Figure 5.**
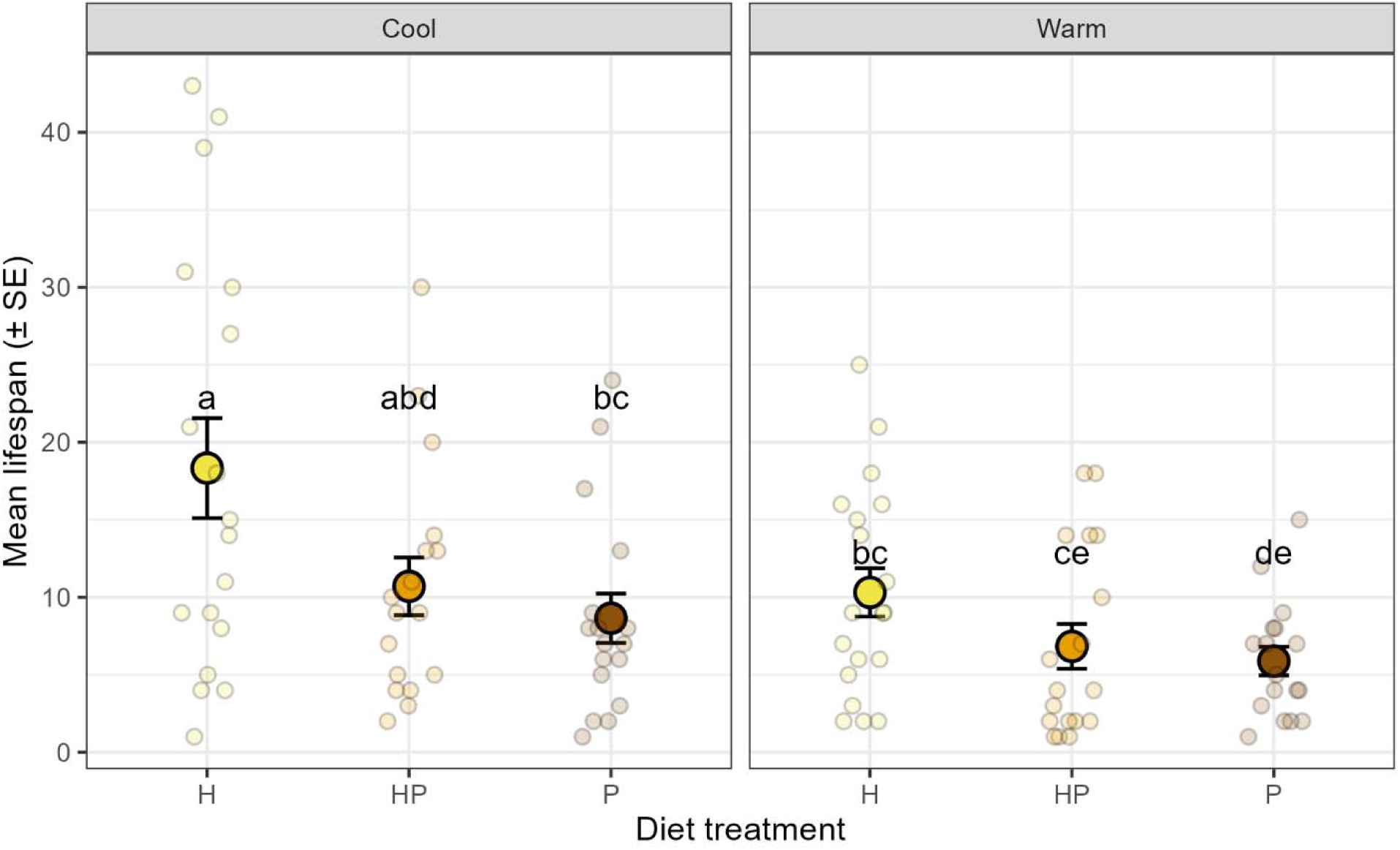
Effect of diet (H – 40% honey; HP – 40% honey and 40% pollen; P – 40% pollen) and temperature treatments (Cool – 16°C day/12°C night; Warm – 22°C day/20°C night) on longevity of adult *Istocheta aldrichi*. Small dots show the lifespan of individual adults; large dots with error bars show estimated marginal means and standard errors. Pairwise comparisons among treatments are based on estimated marginal values of the Cox model linear predictor (log hazard), averaged over sex, with Tukey-adjusted contrasts. Different letters indicate significant differences in hazard ratios among treatments. See Table 4 for detailed statistical information.

## Discussion

Several factors, some of them interacting, were important for the successful overwintering of *I. aldrichi* puparia and the survival of emerging adults. These factors included when the parasitized beetles were collected during the previous season, when parasitoid puparia were placed in overwintering conditions, how long they were kept at cool temperatures, how they were overwintered, how they were warmed up in the spring, and the temperature regime and food provided to adult *I. aldrichi*. Below, each stage of the rearing process is discussed in turn. The discussion concludes with provisional rearing guidelines and what remains to be learned to further improve rearing of *I. aldrichi* for research and biological control releases.

### Overwintering of I. aldrichi puparia

In agreement with Simões & Grenier (1999), Experiment 1 found that *I. aldrichi* adult emergence initially increases with the duration for which puparia are chilled (here, at 5°C), although the proportion emergence plateaued with increasing chilling durations. The time to adult parasitoid emergence decreased with chilling duration, however (up to ∼125 days). Decreasing development time with increasing time in diapause has also been observed in other insects (Tauber & Tauber, 1976; Jones et al., 2023). Thus, some *I. aldrichi* complete their diapause even with relatively short chilling durations, but emergence and development time are optimal after at least 125 days of chilling. This shows that by early to mid-March, they had been in cold conditions long enough to complete their diapause, well before average outdoor soil temperatures in British Columbia, Ontario, or Québec typically rise to values (>10°C) at which their development would progress significantly (P. K. Abram, unpublished analyses). Completion of diapause well before the end of insects’ overwintering period is a well-known phenomenon (Tauber & Tauber, 1976). From a practical standpoint, this means that *I. aldrichi* can be removed earlier than they would emerge under field conditions for experimental laboratory work with minimal effects on survival or development time.

Several possibilities for how to overwinter *I. aldrichi* have been identified: outdoors under the soil, in a protected environment outdoors (e.g., a cellar) or in a growth chamber that mimics outdoor conditions (King & Hallock, 1925; Simões & Grenier, 1999; McDonald & Klein, 2023). A study conducted more than 100 years ago in the Northeastern USA showed that *I. aldrichi* had higher survival rates when overwintered in a cold cellar than when overwintered under the soil (King & Hallock, 1925). Simões & Grenier (1999), who conducted their study in the Azores archipelago where winter temperatures are relatively warm, only succeeded in having *I. aldrichi* emerge when stored indoors in a refrigerator at ∼4°C.

In Experiment 2, there were context-dependent consequences of overwintering puparia under the soil versus indoors in a growth chamber at constant ∼5°C; overwintering conditions interacted with spring conditions and the collection dates of samples the previous summer (see below) to influence the proportion of *I. aldrichi* emerging. Notably, samples from parasitized *P. japonica* collected later in the previous season (i.e., the end of July and beginning of August) seemed to have more greatly reduced *I. aldrichi* emergence when overwintered indoors than outdoors. In contrast, under outdoor spring conditions, fewer adults emerged from puparia overwintered outdoors compared to those overwintered indoors. Overwintering puparia indoors is simpler, but risks growth chamber malfunctions that can cause rapid warming and destruction of samples (which was encountered in subsequent years). Overwintering puparia outdoors, whether under the soil or in a protected structure (e.g., cellar) is not prone to these malfunctions and so may be safer, as long as winter temperatures in a given study region are sufficiently cold (but not too cold) and samples are well protected from wildlife, hyperparasitoids, and predation from soil invertebrates. Indeed, overwintering samples outdoors in a region of Canada (south coastal British Columbia) with relatively mild winters still produced satisfactory results as long as puparia were brought indoors for subsequent adult *I. aldrichi* emergence in the spring. It is stressed, however, that if constant indoor temperatures are used, 5°C is not necessarily the optimal overwintering temperature, and other overwintering temperatures should be tested in future studies.

The follow-up overwintering experiment to test the importance of fall burial date (Experiment 3) clearly showed that if *I. aldrichi* puparia are buried outdoors, it is critical to do so as early as possible once *I. aldrichi* have entered the pupal stage. This might imply that relatively stable cool temperatures, as occur under the soil, between late summer and winter are important for subsequent overwintering survival. Indeed, the earliest burial date produced similar survival to what was observed in Experiment 2, where puparia were kept in stable, cool conditions (15°C) in the fall before overwintering. Likely, exposure to higher temperatures before the overwintering period, as would have occurred in the two later-buried treatments in Experiment 3, results in increased depletion of metabolic resources that are needed to survive the winter (Feder et al. 1997; Hahn and Denlinger 2007). Humidity during this period may also be an important factor. Further studies are needed to more precisely define the range of temperature and humidity conditions that are optimal for *I. aldrichi* puparia between when they form puparia in the summer and the onset of cold temperatures in the winter.

### Rearing *I. aldrichi* puparia for adult emergence in the spring

Previously, it was unclear whether *I. aldrichi* puparia needed to be gradually warmed up after overwintering (as occurs in the soil outdoors), or if they could simply be held at cold temperatures for longer periods and then moved to warmer temperatures to complete their development as needed. In Experiment 2, gradual warming mimicking temperature patterns of soil outdoors did not consistently produce higher emergence levels than keeping samples in 5°C in the spring and then moving them to 23°C in the early summer. Thus, the latter strategy is a simple and feasible option for producing *I. aldrichi* emergence at planned intervals in the spring and summer. Experiment 3 also showed that the duration of post-overwintering puparia storage in cool conditions (12°C) can be varied to stagger *I. aldrichi* emergence over the summer period without affecting their survival.

However, mimicking soil temperatures in growth chambers may be desirable when emerging *I. aldrichi* adults are used for biological control releases, as it would be expected to result in emergence timing that is better synchronized with outdoor *P. japonica* populations, given inter-annual spring temperature variation. Indeed, the emergence timing of the spring treatments that followed outdoor soil temperatures (Experiment 2; treatments I and II) closely matched the beginning of *P. japonica* emergence in British Columbia in 2023, while emergence in treatment III (1.5°C lower than soil temperatures) overlapped with a period of high relative *P. japonica* abundance (Canadian Food Inspection Agency, unpublished data). In subsequent years of *I. aldrichi* releases for biological control of *P. japonica* (2024, 2025), different variations on outdoor spring soil temperatures in incubators were therefore used to produce staggered emergence of *I. aldrichi* that overlapped with periods of high host abundance (Abram, 2026).

The only spring treatment that was clearly inferior in terms of *I. aldrichi* emergence was when they were kept in a plastic tote underground, especially samples that were overwintered indoors. It was noted that humidity was very high inside these poorly ventilated totes and perhaps that may have negatively affected adult emergence. In addition, it was inconvenient to collect emerging *I. aldrichi* adults from a buried container. Thus, it is recommended that substrate containing emerging *I. aldrichi* be moist and held under well-ventilated conditions indoors.

### Feeding and temperature conditions for adult *I. aldrichi*

Prior research on the nutrition of adult tachinids showed that feeding on sugary substances and/or pollen can accelerate egg production, increase survival and extend longevity (Dindo & Grenier, 2014; Dindo et al., 2019), and it has been suggested that *I. aldrichi* may benefit from feeding on nectar from several different plants (McDonald & Klein, 2023). In Experiment 4, adult *I. aldrichi* lived the longest when provided with honey and the shortest when provided with pollen. Somewhat counterintuitively, providing both honey and pollen was inferior to providing honey alone. While pollen could provide a source of proteins and amino acids and has been found to have a longevity-promoting effect in other Tachinidae (Dindo et al., 2019), under this study’s conditions it could have potentially harbored bacteria or fungi that reduced *I. aldrichi* longevity. It is also possible that the specific type of pollen used (apple pollen) is not suitable for *I. aldrichi*, but other pollen types would be. More research is clearly needed to identify which specific nutrients best support *I. aldrichi* longevity, and how nutrition affects fecundity. While fecundity of the *I. aldrichi* adults in the different treatments was not measured, mating was observed in emergence and release cages. In addition, *I. aldrichi* adults parasitized *P. japonica* under field conditions after they were released (Makovetski & Abram, 2023; Abram, 2026), indicating that they were fertile. Not surprisingly, lower temperatures also extended the longevity of *I. aldrichi*; those fed with honey water and kept at cool temperatures lived the longest of any of the treatments and this was the condition subsequently chosen for storing *I. aldrichi* adults between their emergence and release for biological control of *P. japonica*. While the nutritional and storage temperature regime used is not yet optimized, providing adults with honey-water does extend their longevity and can clearly produce fertile individuals that are robust, mated, and ready to parasitize hosts once released in the field.

### Provisional guidelines and future research

Based on the results presented here, the following provisional guidelines for rearing, overwintering, and staggering emergence of *I. aldrichi* are proposed (see Figure 6 for a graphical summary). Parasitized *P. japonica* should be collected as early as possible during the growing season, ideally during the first two weeks during which they are present. If they are collected using traps, plant material and moist substrate (e.g., potting soil) should be placed in the bottom of the trap (Legault et al., 2026). To maximize the size of *I. aldrichi* puparia produced, parasitized beetles should be fed with plant material in ventilated containers containing moist vermiculite in the laboratory for approximately 1 week after which the plant material should be removed (Legault et al., 2026). After *I. aldrichi* from parasitized beetles have developed to puparia, these puparia should be stored in moist vermiculite and placed in conditions with stable, cool temperatures (< 15°C), either under the ground or in a reliable growth chamber as soon as possible, ideally in late August. If being overwintered in a growth chamber, they can be moved to cooler temperatures (∼5°C) in November. If adult *I. aldrichi* are needed for laboratory experiments, puparia can be unearthed (if overwintered outdoors) or removed from the growth chamber and moved to warmer temperatures as early as February. If being used for biological control releases, emergence of *I. aldrichi* in the spring and early summer can be scheduled either by warming them up gradually in incubators set at temperatures based on outdoor soil temperatures, or keeping them in incubators at constant cool conditions (5–12°C) throughout the spring and then moving them to warm conditions (∼22°C) to emerge gradually in the early summer period before releases. As *I. aldrichi* adults emerge, they should be moved to relatively cool conditions (16°C day, 12°C night) and fed with honey-water until they are released.

**Figure 6.**
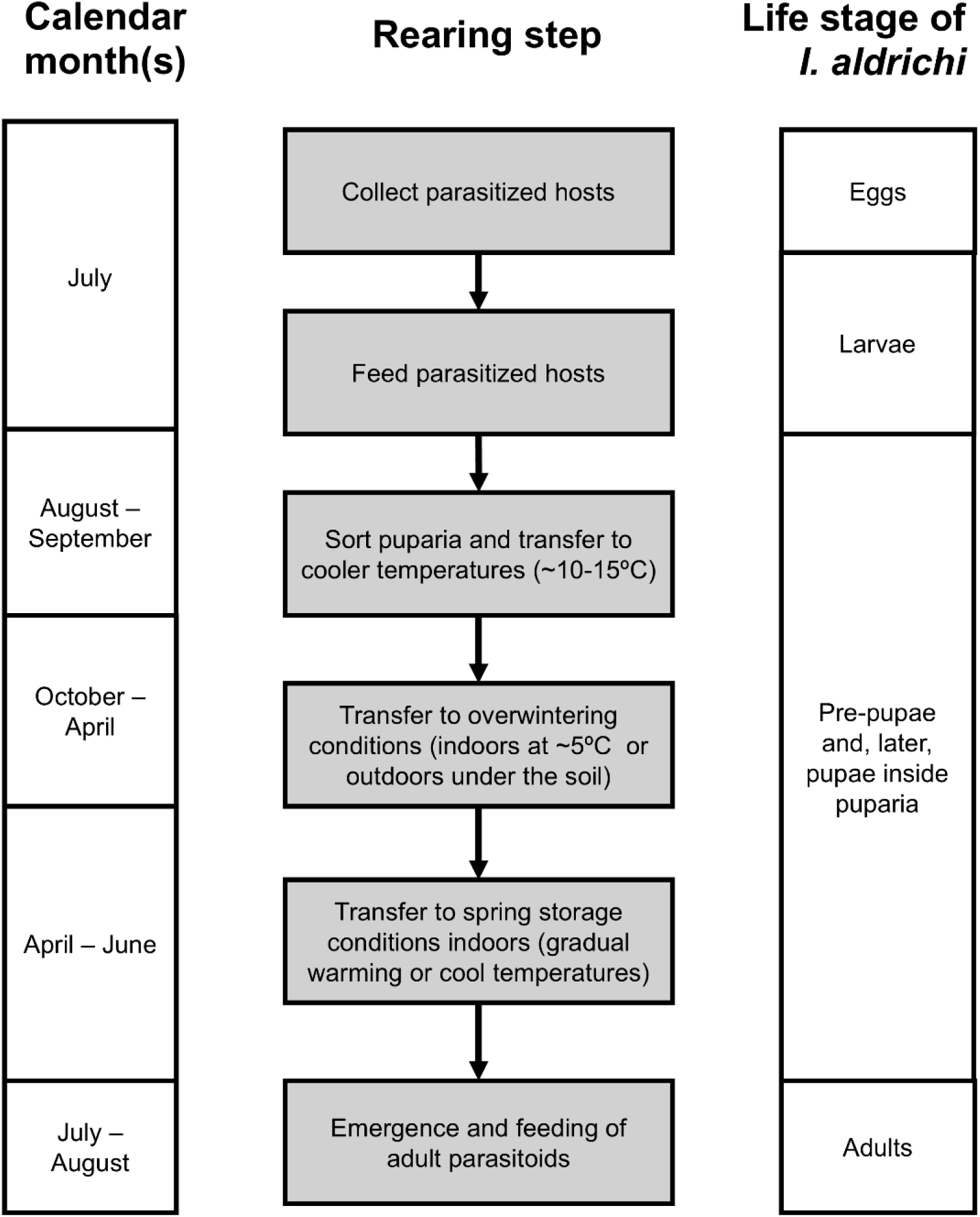
Overview of the steps to rear the parasitoid *Istocheta aldrichi* from field-collected *Popillia japonica* hosts, with corresponding calendar months and life stages (for Ontario and Québec, Canada). Note that the exact timing of the formation of pre-pupae and pupae with respect to the rearing steps and calendar months is not known.

Although several factors important for *I. aldrichi* rearing were identified, substantial knowledge gaps remain. Notably, the overwintering and spring conditions tested produced relatively low average emergence levels (<50%) indicating a need to further improve on these conditions. While there is ample potential for strategically timing and staggering *I. aldrichi* emergence with different spring temperature regimes, development of a predictive degree-day model would greatly refine the planning and implementation of these regimes (L. M. Seehausen, personal communication). Finally, there is a need to determine the importance of humidity conditions at different stages of *I. aldrichi* development, and refine nutritional regimens for adult *I. aldrichi* to reduce the observed variation in longevity. As interest in *I. aldrichi* as a biological control agent of *P. japonica* has recently resurged, the findings reported here and in the companion study (Legault et al., 2026), provide an important foundation for improving rearing protocols, understanding the parasitoid’s biology, and supporting its use in suppressing *P. japonica* populations as the pest continues to expand its global range.

## Acknowledgements

We thank Jessie Moon, Fina VanderPloeg, Warren Wong, Clarissa Capko, Jiamin Bae, Yonathan Uriel, Michelle Franklin, Liam Duff, Juliette Matsukubo, Laura Kostyniuk, Josiane Vaillancourt, Gaétan Racette, Natali Demers, Angélie Laplante, Marianne Larose, Laurence Bélanger-LaFaille, Sabrina Tardy, and Julien Robert for help in the field and laboratory. We thank Peter Mason and Lauren Henry for providing helpful suggestions on earlier drafts of the manuscript. This project was supported by funding from Agriculture and Agri-Food Canada (Projects J-003401, J-002839, J-002201) and the Natural Sciences and Engineering Research of Canada (NSERC) to JB. We thank James O’Hara for taxonomic support and Richard McDonald for helpful advice on *I. aldrichi* rearing provided through e-mail correspondence at the beginning of the project.

## References

Abram PK (2026) Istocheta aldrichi (Mesnil), a biological control agent of the Japanese beetle, Popillia japonica Newman, establishes in British Columbia. The Tachinid Times (in press).

Allen GR & Hunt J (2001) Larval competition, adult fitness, and reproductive strategies in the acoustically orienting ormiine *Homotrixa alleni* (Diptera: Tachinidae). Journal of Insect Behavior 14: 283–297.

Barr DJ, Levy R, Scheepers C & Tily HJ (2013) Random effects structure for confirmatory hypothesis testing: keep it maximal. Journal of Memory and Language 68: 255–278.

Bates D, Mächler M, Bolker B & Walker S (2015) Fitting linear mixed-effects models using lme4. Journal of Statistical Software 67: 1–48.

Brodeur J, Doyon J, Abram PK & Parent JP (2024) Popillia japonica Newman, Japanese beetle/scarabée japonais (Coleoptera: Scarabaeidae). In: Vankosky MA & Martel V (eds) Biological Control Programmes in Canada 2013–2023. CABI, Wallingford, UK, pp. 343–350.

Canadian Food Inspection Agency (CFIA) (2025) British Columbia Japanese beetle survey reports. Available at: https://inspection.canada.ca/en/plant-health/invasive-pests-and-plants/insects/japanese-beetle/japanese-beetle-bc

CABI (2021) Classical biological control of Japanese beetle. Available online at: https://www.cabi.org/projects/classical-biological-control-of-japanese-beetle/

Cingolani MF, Barakat MC, Cerretti P, Chirinos DT, Ferrer F, Gaviria Vega J, Grenier S et al. (2025) Dipteran parasitoids as biocontrol agents. BioControl 70: 1–16.

Clausen CP, King JL & Teranishi C (1927) The parasites of Popillia japonica in Japan and Chosen (Korea), and their introduction into the United States. US Department of Agriculture Bulletin 1429.

Dindo ML & Grenier S (2014) Production of dipteran parasitoids. In: Morales-Ramos JA, Rojas MG & Shapiro-Ilan DI (eds) Mass Production of Beneficial Organisms: Invertebrates and Entomopathogens. Academic Press, London, UK, pp. 101–143.

Dindo ML, Rezaei M & De Clercq P (2019) Improvements in the rearing of the tachinid parasitoid *Exorista larvarum* (Diptera: Tachinidae): influence of adult food on female longevity and reproductive capacity. Journal of Insect Science 19: Article 6.

Feder JL, Stolz U, Lewis KM, Perry W, Roethele JB & Rogers A (1997) The effects of winter length on the genetics of apple and hawthorn races of *Rhagoletis pomonella* (Diptera: Tephritidae). Evolution 51: 1862–1876.

Fleming WE (1968) Biological Control of the Japanese Beetle. USDA Technical Bulletin 1383, 83 pp.

Gagnon ME, Doyon J, Legault S & Brodeur J (2023) The establishment of the association between the Japanese beetle (*Popillia japonica*) and the parasitoid *Istocheta aldrichi* (Diptera: Tachinidae) in Québec, Canada. The Canadian Entomologist 155: e32.

Hahn DA & Denlinger DL (2007) Meeting the energetic demands of insect diapause: nutrient storage and utilization. Journal of Insect Physiology 53: 760–773.

Hartig F (2024) DHARMa: Residual diagnostics for hierarchical (multi-level / mixed) regression models. R package version 0.4.7, https://CRAN.R-project.org/package=DHARMa.

Hutchinson WD (2023) Long-term biological control of Japanese beetle? A winsome fly update. Available at: https://blog-fruit-vegetable-ipm.extension.umn.edu/2023/08/long-term-biological-control-of.html

Hutchinson WD, Buchholz EK, Meys EL & Wold-Burkness S (2024) The winsome fly, Istocheta aldrichi: a unique biocontrol agent for Japanese beetle in Minnesota. Fruitedge. Available at: https://fruitedge.umn.edu/winsome-fly-istocheta-aldrichi-unique-biocontrol-agent-japanese-beetle-minnesota

Jones IM, Seehausen ML, Smith SM & Bourchier RS (2023) The effects of warm and cold periods on resource depletion and emergence synchrony in diapausing *Hypena opulenta*: implications for biological control of invasive swallow-worts in North America. Entomologia Experimentalis et Applicata 171: 990–997.

King JL & Hallock HC (1925) A report on certain parasites of *Popillia japonica* Newman. Journal of Economic Entomology 18: 351–356.

Lasnier J, de Coussergues CH, Baril A & Vincent C (2025) Abundance of Japanese beetle adults and its parasitoid *Istocheta aldrichi* in a Québec commercial vineyard. Bulletin of Insectology 78: 1–10.

Legault S, Doyon J & Brodeur J (2024) Reliability of a commercial trap to estimate population parameters of Japanese beetles, *Popillia japonica*, and parasitism by *Istocheta aldrichi*. Journal of Pest Science 97: 575–583.

Legault S, Doyon J, Abram PK & Brodeur J (2026). Rearing Istocheta aldrichi (Diptera: Tachinidae) from field-collected Japanese beetles (Popillia japonica): 1. Methods to improve insect collection and parasitoid pupariation. bioRxiv 10.64898/2026.02.18.706618

Lenth R (2023) emmeans: Estimated marginal means, aka least-squares means. R package version 1.8.5.

Ludwig, D., & Fox, H. (1938). Growth and survival of Japanese beetle larvae reared in different media. Annals of the Entomological Society of America, 31(4), 445–456.

Makovetski V & Abram PK (2024) *Istocheta aldrichi* (Mesnil) makes its biological control debut in British Columbia, Canada. The Tachinid Times 37: 4–10.

Makovetski V, Smith AB & Abram PK (2025) Crowdsourced online data as evidence of absence of non-target attack from the century-old introduction of *Istocheta aldrichi* for biological control of *Popillia japonica* in North America. Journal of Pest Science 98: 1451–1462.

McDonald RC & Klein MG (2023) Establishing the winsome fly, *Istocheta aldrichi* Mesnil, a major natural enemy of Japanese beetle, Popillia japonica Newman adults. Available at: 10.13140/RG.2.2.22208.30720.

Nakamura S (1995) Optimal clutch size for maximizing reproductive success in a parasitoid fly, *Exorista japonica* (Diptera: Tachinidae). Applied Entomology and Zoology 30: 425–431.

O’Hara JE (2014) New tachinid records for the United States and Canada. The Tachinid Times 27: 34–40. Available at: http://www.nadsdiptera.org/Tach/WorldTachs/TTimes/Tach27.html

Pelletier M, Legault S, Doyon J & Brodeur J (2023) Where and why do females of the parasitic fly *Istocheta aldrichi* lay their eggs on the body of adult Japanese beetles? Journal of Insect Behavior 36: 308–317.

Potter DA & Held DW (2002) Biology and management of the Japanese beetle. Annual Review of Entomology 47: 175–205.

R Core Team (2024) R: A language and environment for statistical computing. R Foundation for Statistical Computing, Vienna, Austria. Available at: https://www.R-project.org/

Ritz C, Baty F, Streibig JC & Gerhard D (2015) Dose–response analysis using R. PLOS ONE 10: e0146021.

Shanovich HN, Dean AN, Koch RL & Hodgson EW (2019) Biology and management of Japanese beetle (Coleoptera: Scarabaeidae) in corn and soybean. Journal of Integrated Pest Management 10: 1–14.

Simões AM & Grenier S (1999) An investigation into the overwintering capability of Istocheta aldrichi (Mesnil) (Diptera: Tachinidae), a parasitoid of Popillia japonica Newman (Coleoptera: Scarabaeidae) on Terceira Island, Azores. Universidade dos Açores. Available at: http://hdl.handle.net/10400.3/140.

Stilwell PA, Culotta JA, Hutchison WD & Lindsey AR (2025) The genome of *Istocheta aldrichi* (Diptera: Tachinidae), a parasitoid of the Japanese beetle, Popillia japonica (Coleoptera: Scarabaeidae). bioRxiv 2025–11. 10.1101/2025.11.02.686142

Stireman JO III, O’Hara JE & Wood DM (2006) Tachinidae: evolution, behavior, and ecology. Annual Review of Entomology 51: 525–555.

Strangi A, Paoli F, Nardi F, Shimizu K, Kimoto T, Iovinella I, Bosio G, Roversi PF, Carapelli A & Marianelli L (2024) Tracing the dispersal route of the invasive Japanese beetle *Popillia japonica*. Journal of Pest Science 97: 613–629.

Tauber MJ & Tauber CA (1976) Diapause maintenance, termination, and postdiapause development. Annual Review of Entomology 21: 81–107.

