## Supplementary Material for "Rearing *Istocheta aldrichi* (Diptera: Tachinidae) from field-collected Japanese beetle (*Popillia japonica*): 2. Methods to improve overwintering, adult emergence and longevity"

[1] Agriculture and Agri-Food Canada, Agassiz Research and Development Centre, Agassiz, BC, Canada; [2] Université de Montréal, Institut de recherche en biologie végétale, Département de sciences biologiques, Montréal, QC, Canada; [3] University of Victoria, Department of Biology, Victoria, BC, Canada; [4] Agriculture and Agri-Food Canada, Ottawa Research and Development Centre, Ottawa, ON, Canada; [5] Agriculture and Agri-Food Canada, Saint-Jean-sur-Richelieu Research and Development Centre, Saint-Jean-sur-Richelieu, QC, Canada

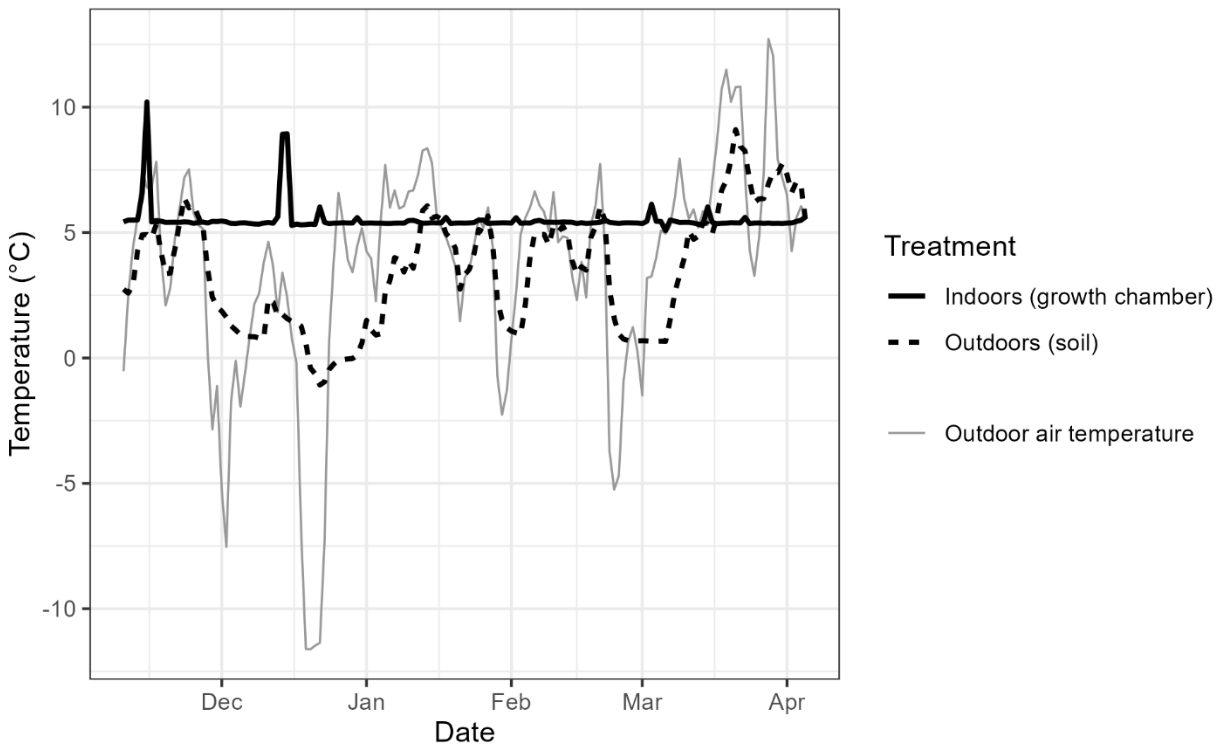


**Figure S1.** The average daily air temperature of the two different *Istocheta aldrichi* puparia overwintering treatments over time in 2022/2023 for Experiment 2. Outdoor average daily air temperature is also shown, for reference.

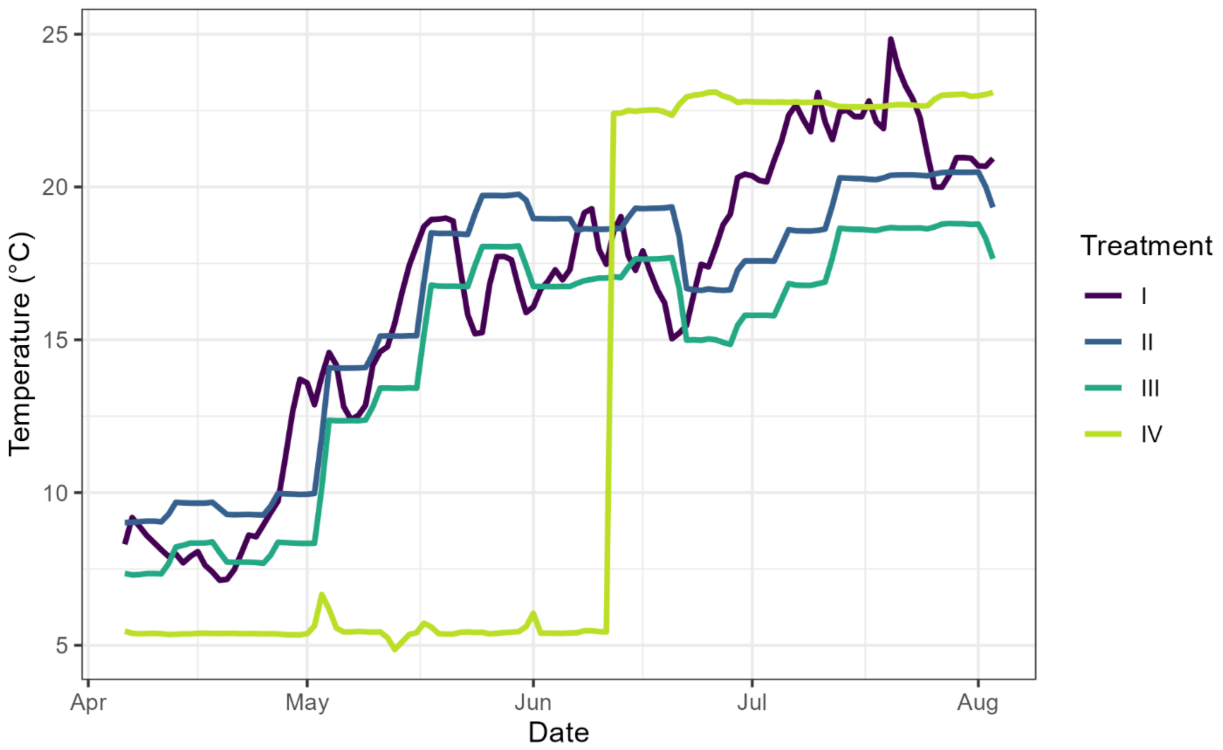


**Figure S2.** The average daily air temperature of different *I. aldrichi* puparia spring treatments over time in 2023 for Experiment 2. I – outdoors under the soil; II – indoors at soil temperatures; III – indoors at lower-than-soil temperatures; IV – indoors at 5°C and then moved to ~22°C.
